# Amplitude and phase coupling optimize information transfer between brain networks that function at criticality

**DOI:** 10.1101/2021.03.15.435461

**Authors:** Arthur-Ervin Avramiea, Anas Masood, Huibert D. Mansvelder, Klaus Linkenkaer-Hansen

**Author notes:** Corresponding author: Klaus Linkenkaer-Hansen. Department of Integrative Neurophysiology, Center for Neurogenomics and Cognitive Research, (CNCR), VU University Amsterdam, 1081 HV Amsterdam, Netherlands.

## Abstract

Brain function depends on segregation and integration of information processing in brain networks often separated by long-range anatomical connections. Neuronal oscillations orchestrate such distributed processing through transient amplitude and phase coupling, yet surprisingly little is known about local network properties facilitating these functional connections. Here, we test whether criticality—a dynamical state characterized by scale-free oscillations—optimizes the capacity of neuronal networks to couple through amplitude or phase, and transfer information. We coupled *in silico* networks with varying excitatory and inhibitory connectivity, and found that phase coupling emerges at criticality, and that amplitude coupling, as well as information transfer, are maximal when networks are critical. Importantly, regulating criticality through neuromodulation of synaptic strength showed that critical dynamics—as opposed to a static ratio of excitatory and inhibitory connections—optimize network coupling and information transfer. Our data support the idea that criticality is important for local and global information processing and may help explain why brain disorders characterized by local alterations in criticality also exhibit impaired long-range synchrony, even prior to degeneration of axonal connections.

## Introduction

Segregation of function across cortical areas allows for economical wiring and reduces metabolic costs (Achard and Bullmore, 2007; Sporns, 2010; Wang and Clandinin, 2016). However, to function adaptively in a dynamic environment, the brain also has to flexibly integrate its activity across distant cortical regions. Neuronal oscillations are an important network-level mechanism driving coordination not only in local cortical assemblies but also through amplitude and phase coupling of distant brain regions (Varela et al., 2001; Fries, 2005). In fact, mapping functional connectivity in the brain often relies on quantifying such phase (Sauseng et al., 2005; Stam et al., 2007; Palva et al., 2010; Crespo-Garcia et al., 2013; Marzetti et al., 2019; Tewarie et al., 2019) and amplitude dependencies (Daffertshofer and van Wijk, 2011; Hipp et al., 2012; Vidal et al., 2012; Foster et al., 2015; O’Neill et al., 2015; Tewarie et al., 2019). However, although coupling of oscillations enables information exchange, a fully synchronized brain is also not advantageous, since such a stable state does not allow for creation of new information (Tagliazucchi and Chialvo, 2013; Tognoli and Kelso, 2014). For this reason, several investigators have argued that the brain resides optimally in a state that is neither fully segregated nor fully integrated, where transient changes in phase and amplitude coupling allow it to explore the largest repertoire of functional states supported by the static structural connectome (Tagliazucchi and Chialvo, 2013; Deco and Kringelbach, 2016; Li et al., 2019). The so-called “critical state” has often been suggested as a statistically well-defined dynamical state that could fulfill the contrasting demands of stability and susceptibility to inputs (Bak et al., 1987; Chialvo, 2010). Indeed, the hallmark of the critical state is spatiotemporal dynamics with power-law scaling behavior (Linkenkaer-Hansen et al., 2001; Beggs and Plenz, 2003), indicating that spatial and temporal patterns on a wide range of scales can occur.

In a computational model of critical oscillations (CROS) (Poil et al., 2012; Dalla Porta and Copelli, 2019; Avramiea et al., 2020), it was shown that for any level of inhibitory connectivity, scale-free fluctuations in the amplitude of oscillations appeared when a suitable amount of excitatory connections were added to the network, indicating that critical dynamics of oscillations requires a specific balance of excitation and inhibition. Interestingly, at the same balance we find scale-free distributions of neuronal avalanches—another hallmark of critical brain dynamics (Beggs and Plenz, 2003). Both neuronal avalanches (Kinouchi and Copelli, 2006; Shew et al., 2009; Gautam et al., 2015) and critical oscillations (Avramiea et al., 2020) have been associated with the widest dynamic range to stimuli. This suggests that at the critical point, networks might be most responsive to inputs coming from distant brain areas. Consequently, alterations in the local criticality of networks might lead to impaired coupling. In agreement with this idea, brain disorders such as Alzheimer’s (Montez et al., 2009) and schizophrenia (Nikulin et al., 2012) have been associated with alterations in criticality, excitation/inhibition balance (Lisman, 2012; Busche and Konnerth, 2016), and reduced long-range phase coupling in fronto-parietal networks (Babiloni et al., 2004; Sheffield et al., 2015). Together, these findings suggest that criticality, excitation/inhibition balance, and long-range coupling might be interdependent.

Here, we hypothesize that local criticality of neuronal networks is required for establishing long-range coupling in the amplitude and phase of neuronal network oscillations, as well as for transferring information. We tested this hypothesis in the CROS model (Poil et al., 2012; Dalla Porta and Copelli, 2019; Avramiea et al., 2020). We connected two networks and varied their E/I ratio and, consequently, the level of criticality, to understand its impact on long-range coupling and information transfer. We found that information transfer is optimized in the critical regime, where networks show maximal coupling in the phase and amplitude of oscillations. Thus, regulation of criticality may be important for long-range communication, and alterations in criticality may explain disruptions in long-range synchrony in brain disorders, along with their functional consequences.

## Results

### Multi-level criticality is preserved after coupling networks

To model communication between two distant brain areas, e.g., frontal and parietal, we created two networks of excitatory and inhibitory neurons, and coupled them through *N* long-range connections (Fig. 1A). We first selected at random *N/2* excitatory positions from the first network and created synapses to excitatory neurons at the same positions in the second network, and repeated the operation for the second network, making sure that the reciprocal connections did not run through the same two neurons. We set a conduction delay of 25 ms across these long-range connections, comparable with delays across long-range myelinated axons in the brain (Ringo et al., 1994). We controlled local E/I balance by changing the percentage of local neighbors that each excitatory and inhibitory neuron connects to. It has previously been shown that depending on this E/I balance, multi-level critical behavior emerges in the form of long-range temporal correlations (LRTC) in the amplitude modulation of oscillations and power-law scaling of neuronal avalanche sizes (κ) (Poil et al., 2012). When the networks are not connected (*N* = 0), we see that LRTC (as measured by the detrended fluctuation analysis exponent, DFA) and scale-free avalanches do indeed occur for the same ratio of excitatory and inhibitory synapses (Fig. 1B,C). After coupling networks with the same local E/I ratio, we compared the DFA and κ index of avalanches, and found them to be similar between the two networks (Fig. 1D,E). This allows us to quantify criticality of the coupled networks as the average of the DFA exponents or κ indices computed separately for the two networks. We find that, after coupling the two networks, the critical line shifts down in the phase space: with the additional input from the other network, each network requires less local excitatory connections to achieve critical fluctuations (Fig. 1F,G). Importantly, the relationship between LRTC and avalanches is preserved after coupling the networks, and depends on a specific balance between excitation and inhibition (Fig. 1F,G).

**Figure 1.**
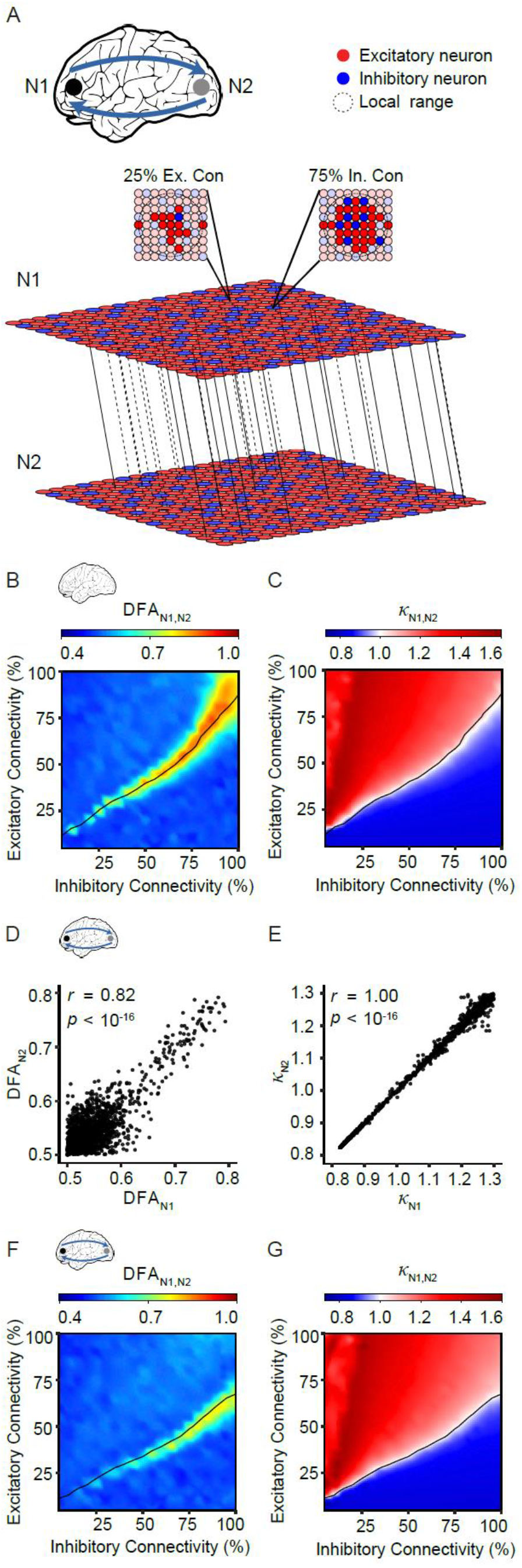
Coupled networks exhibit critical dynamics in the form of long-range temporal correlations and avalanches with power-law scaling. **A)** We coupled two networks consisting of excitatory and inhibitory neurons arranged on a grid. Local connectivity is set separately for excitatory and inhibitory neurons as the percentage of neurons within a local range of 4 neurons that each neuron connects to. Long-range connections are defined between excitatory neurons on the two grids: *continuous lines* represent connections from the top to the bottom network, *dashed lines* represent connections from the bottom to the top network. At a certain balance between excitation and inhibition, the unconnected networks exhibit (**B**) long-range temporal correlations (DFA > 0.5), and (**C**) power-law scaling of avalanche sizes (indicated with the *black line*, also in **B**). After connecting the two networks (*N* = 400 long-range connections), at a conduction delay of 25 ms between networks, we find that the two networks have similar criticality, both in terms of long-range temporal correlations **(D)**, and power-law scaling of avalanches **(E)**. Thus, we can use the average DFA exponent or κ to index criticality of the coupled networks. Note that the region in the phase space where the coupled networks exhibit critical dynamics shifts down **(F, G)** compared to the uncoupled networks **(B, C)**.

### Networks exhibit robust amplitude coupling only in the critical regime

To understand the role of criticality for the ability of mutually connected networks to entrain each other, we first investigated amplitude fluctuations of the coupled networks. Co-modulation of critical oscillations was conspicuous at long time scales (Fig. 2A,B). This was confirmed by low-pass filtering the amplitude envelope of the oscillations causing cross-correlations to increase (Fig. 2C). This observation is in line with empirical reports on oscillations in homologous areas (Nikouline et al., 2001; Hipp et al., 2012; Siems and Siegel, 2020), suggesting that long-time-scale co-modulation may emerge from mutual entrainment and does not require a slow drive such as neuromodulation. Interestingly, the capacity of the networks to couple their amplitude fluctuations breaks down both in sub- and super-critical regimes (Fig. 2C–F). To verify that amplitude coupling emerges from a causal interaction between the two networks, we swapped the two halves of one of the signals, which entirely abolished the amplitude correlations for all networks (Fig. 2E,F). The relationship between criticality and amplitude coupling is remarkably robust to changes in the number of connections between the two networks (Fig. S1A) or in conduction delay (Fig. S1B), suggesting that criticality facilitates amplitude entrainment between brain regions at many levels of the cortical hierarchy. When all excitatory neurons in the two networks are coupled to each other, some super-critical networks can also establish amplitude coupling (Fig. S1A); however, this may not be a physiologically realistic scenario and with such a strong integration of the two networks, they may be perceived as one. To assess whether networks with different levels of criticality can also establish amplitude coupling, we allowed for the E/I ratio to vary between the two networks in the ensemble. We found that amplitude coupling emerged only when both networks were close to the critical regime (Fig. S1C). Thus, local criticality of brain areas appears fundamentally important for establishing amplitude coupling.

**Figure 2.**
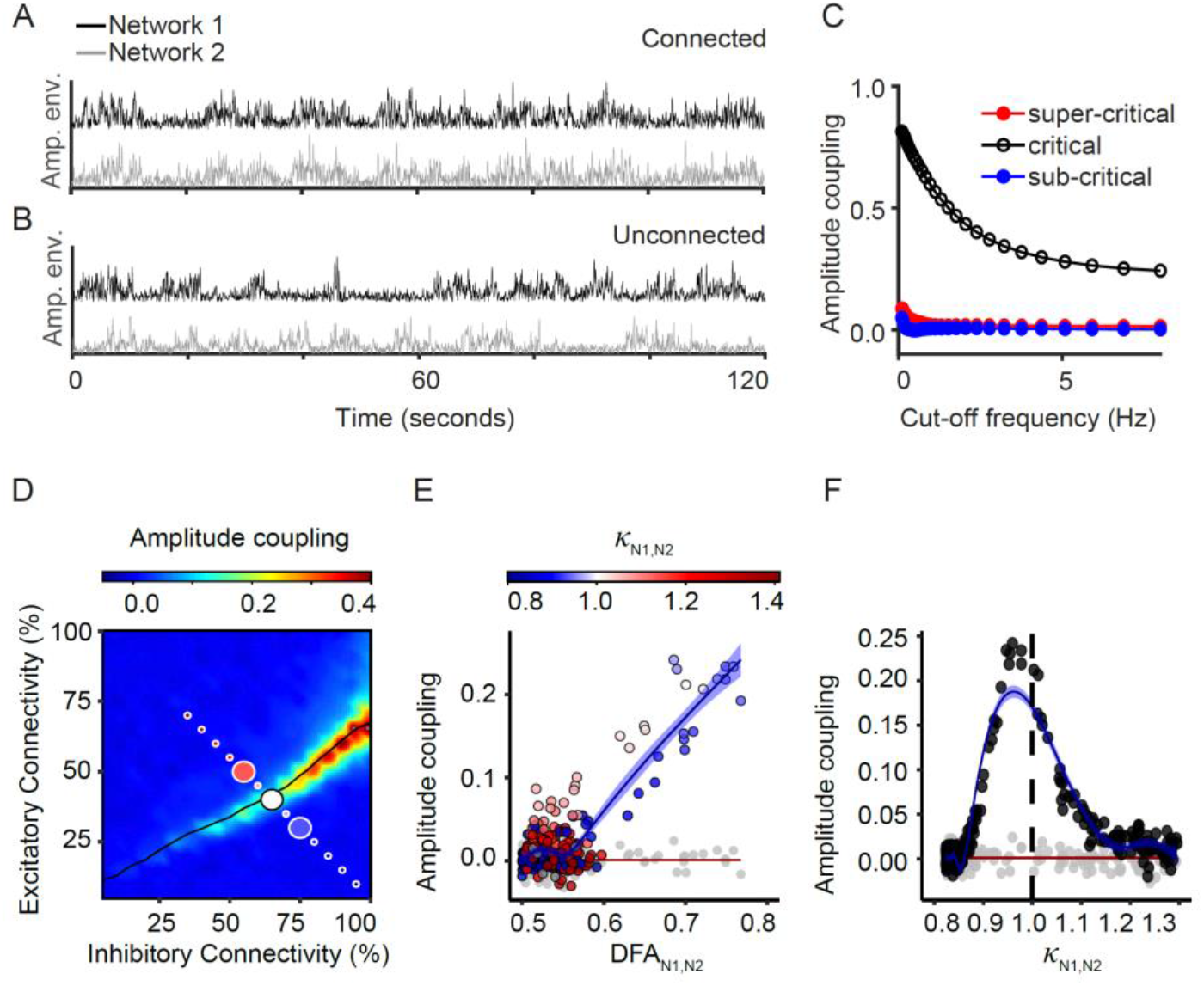
Coupled networks show maximal amplitude correlation at criticality. Two networks exhibiting critical dynamics when connected with *N* = 400 long-range connections and a 25 ms conduction delay, show a striking co-modulation of amplitude visualized at a long time-scale **(A)** whereas no such relationship is seen for unconnected networks **(B)**. **(C)** Lowering the low-pass filter cut-off applied to the amplitude envelope of alpha-band oscillations reveals stronger amplitude co-modulation at low frequencies, albeit only for critical networks. The three lines reflect mean correlation coefficients computed across 5 runs for each of the networks indicated with the corresponding color in the phase space in (**D**). **(D)** We coupled two networks with the same local excitatory and inhibitory connectivity, and ran simulations for all combinations of excitatory/inhibitory connectivity. We found that the correlation of amplitude envelopes peaks in the critical regime, where excitation and inhibition are in balance (*black line* corresponds to κ =1). To further analyze the relationship between amplitude correlation and criticality we sampled networks with excitatory/inhibitory local connectivity from a diagonal crossing the critical line (**D**). The correlation of the amplitude envelopes of the two connected networks peaks in the critical state as measured both with DFA (**E**, *blue line*) and κ (**F**, *blue line*). Surrogates generated by swapping the two halves of the second network before computing the amplitude envelope, show no such relationship with criticality (**E**, **F**, *gray dots* and *red line*).

### Phase coupling emerges at criticality and extends into (slightly) super-critical regimes

The ability of mutually connected networks to phase-couple is considered important for information exchange (Fries, 2005; Palmigiano et al., 2017). We assessed the extent of phase coupling, by computing the inter-site phase clustering between oscillations in the two networks (Fig. 3A,B). We found that phase coupling emerges at criticality and further increases when moving into the super-critical regime (Fig. 3B-E). To verify that phase coupling emerges from a causal interaction between the two networks, we swapped the two halves of one of the signals, which entirely abolished the phase coupling between the two signals (Fig. 3D,E). Next, we varied the number of long-range connections and observed that the relationship between the level of criticality and phase coupling is robust to increases in the number of long-range connections (Fig. S2A). Varying the conductance delays, however, revealed a clear preference to phase couple in certain ranges, consistent with a coupling in-phase or out-of-phase with alpha oscillations (Fig. S2B). Last, we assessed whether networks with different levels of criticality can also establish phase coupling. Varying the E/I ratio in the two networks independently of each other, we found that phase coupling is particularly strong when a super-critical network is connected to a critical one (Fig. S2C).

**Figure 3.**
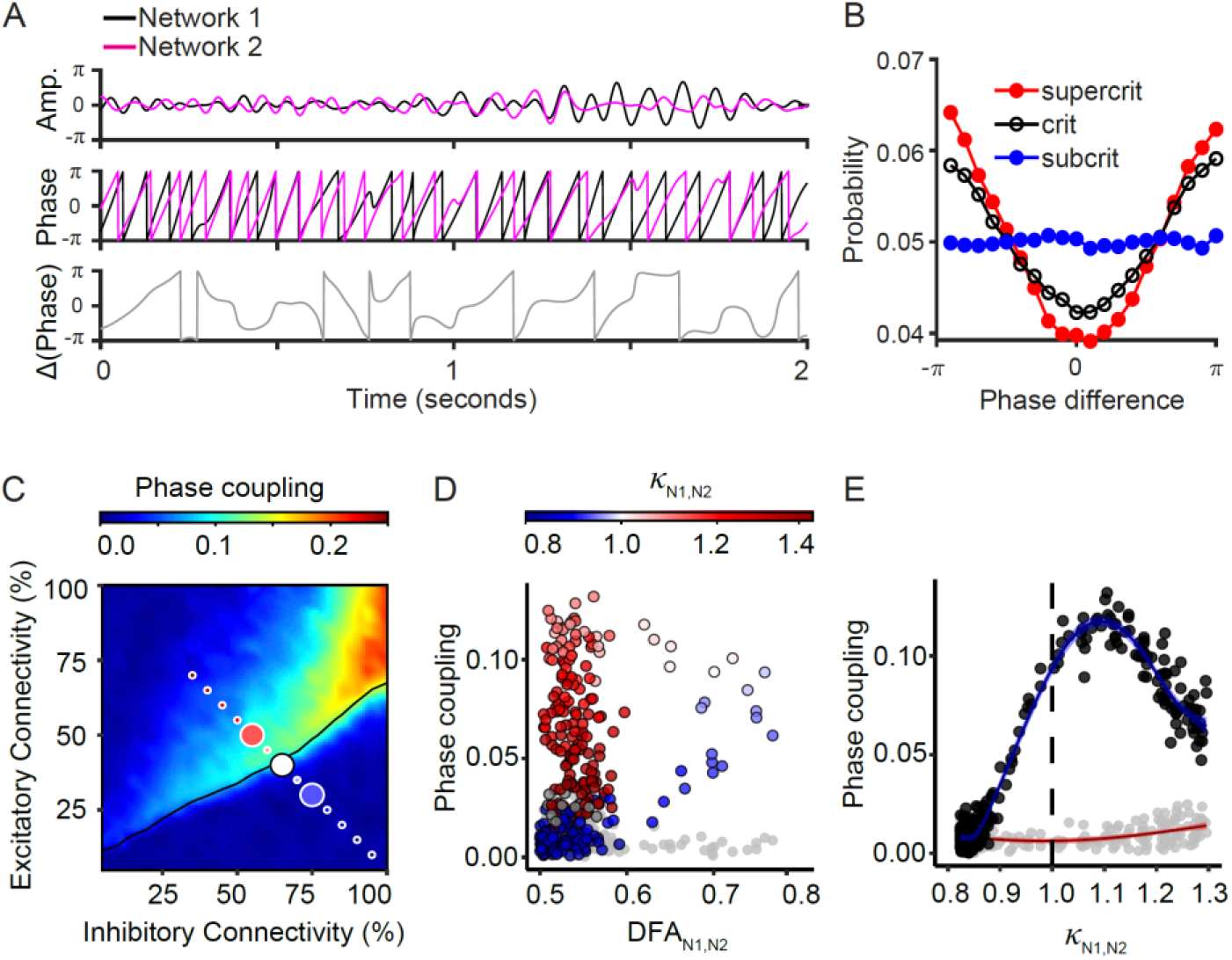
Phase coupling emerges at criticality. **(A)** We connected two networks with *N* = 400 long-range connections each way and a 25 ms conduction delay between networks. We filtered the two signals in the alpha band (*top*), extracted the phase of each signal from the angle of the Hilbert transform (*middle*), and computed the phase difference (*bottom*). **(B, C)** To determine the level of phase coupling between the networks, we computed inter-site phase clustering (ISPC). Phase differences are binned between −π and π, and a phase coupling of 0 implies a uniform distribution of phase differences across bins, whereas a phase coupling of 1 would mean that all phase differences cluster in the same bin. The three lines reflect average phase distributions computed across 5 runs. For each of the three networks we indicated the corresponding color in the phase space of ISPC as a function of excitatory/inhibitory connectivity (**C**). The level of phase clustering depends on criticality: a range of super-critical networks have the highest phase. To further analyze the relationship between phase coupling and criticality, we sampled networks with excitatory/inhibitory local connectivity from a diagonal crossing the critical line (**C**). **(D,E)** Critical networks exhibit an intermediate value of phase coupling (*blue line*). Surrogates generated by swapping the two halves of the second network before computing the ISPC, show no such relationship with criticality (*red line*).

### Information transfer is maximized close to criticality, when networks show maximal amplitude and phase coupling

To ensure that amplitude and phase coupling are not merely epiphenomena of the critical state, but actually enable networks to transfer information over long-range connections, we applied an external stimulus to the first network (Fig. 4A). The input stimulus was Gaussian white noise convolved with an exponential with a time constant τ = 50 ms (Mainen and Sejnowski, 1995). This convolution slows down the stimulus, allowing the networks to keep up with changes in the signal (Fig. 4B). We then correlated the input signal with the amplitude envelope of the two networks. We found that the input is transferred to both networks (Fig. 4C). As expected, the signal takes longer to reach the second network, and some information is lost across networks. As a measure of information transfer, we used the maximal possible correlation (across time lags) between the input signal and the amplitude envelope of oscillations in the second network. Although the external input makes the first network more super-critical in its activity dynamics (Fig. 4D), information transfer is maximized when the second, coupled network, is in a regime close to criticality (Fig. 4E), where amplitude coupling (Fig. 4F) and phase coupling (Fig. 4G) between the two networks are maximized. To verify that information transfer emerges from a causal interaction between the input and the two networks, we swapped the two halves of the amplitude envelopes, which entirely abolished information transfer (Fig. 4E-G). We found that our networks can track inputs better when they are slower (Fig. S3A), and when there are more long-range connections (Fig. S3B). Longer conduction delays over the long-range connections increase the amount of time that it takes the input to reach the second network (Fig. S3C); however, the peak of the correlation of the input with the second network is unaffected (Fig. S3D). Altogether, regardless of the parameter manipulation, information transfer is maximized when the second network is in a critical regime, where phase and amplitude coupling are strongest.

**Figure 4.**
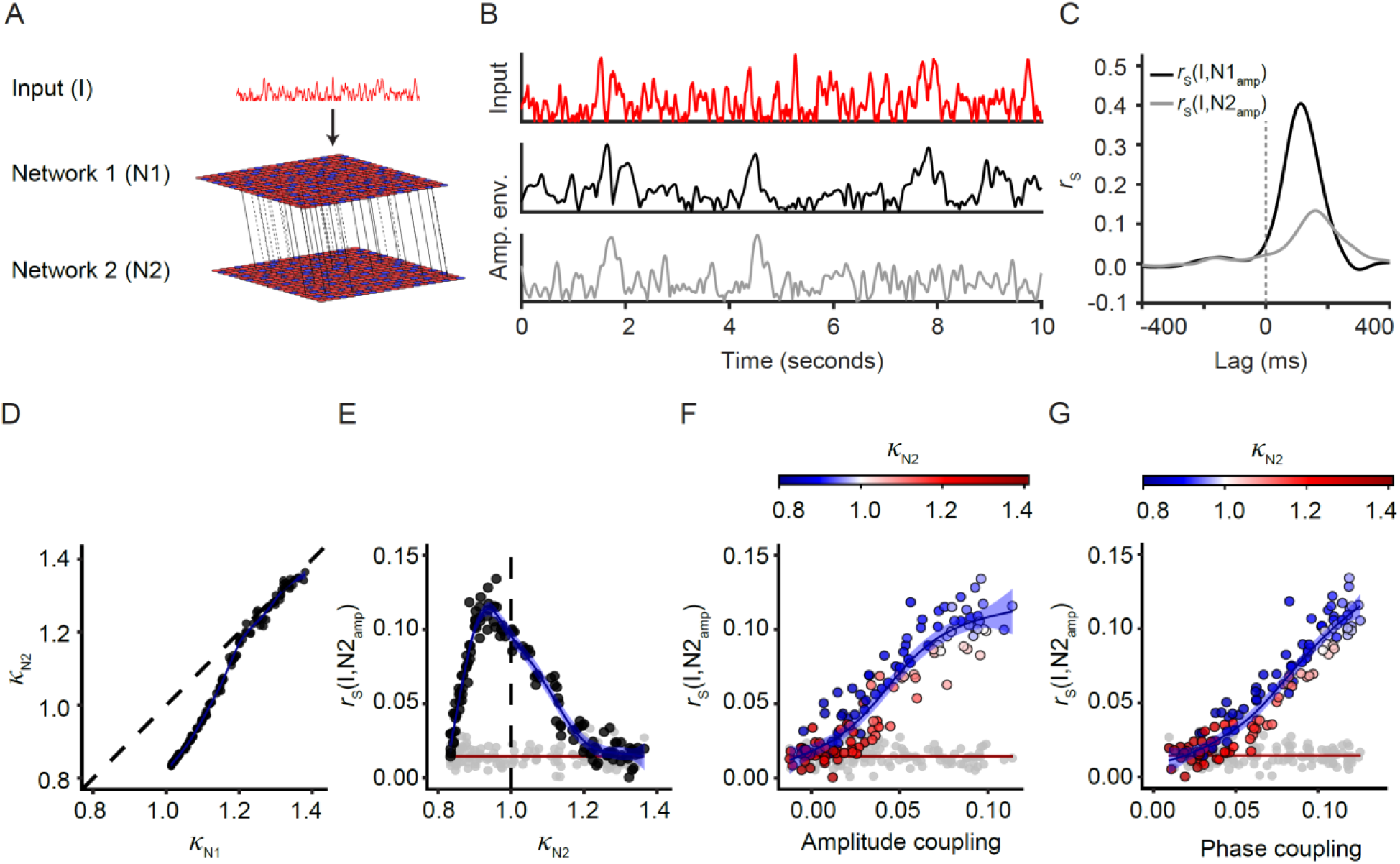
Near-critical networks jointly optimize stimulus-information transfer and amplitude and phase coupling. (**A**) All excitatory neurons in the first network are driven by a white noise signal which was convolved with an exponential with a timescale τ_*input*_= 50 ms. Networks were connected with *N*=400 connections with conduction delay of 25 ms. (**B**) The input (*red*) signal is reflected in the amplitude envelope of the receiving network (*black*). By the time it reaches the second network (*grey*), some information about the input is lost. (**C**) Correlation between the input and the amplitude envelope of the two networks shows that the second network (N2) receives the stimulus ~50 ms later than the first network (N1). We used the highest correlation coefficient of the stimulus with the second network (across time lags) as an indicator of information transfer. **(D**) We sampled pairs of networks with the same ratio of local excitatory/inhibitory connections from the diagonal crossing the critical line, in Fig. 2B. The external input to N1 drives its activity to become more super-critical than N2, as reflected in the right-shift of the κ values relative to the dashed line. We measured the transfer of information from the input signal to N2, as the correlation of the stimulus with the amplitude envelope of the alpha oscillations in N2. Stimulus coupling peaks when N2 is close to critical (**E**), where the amplitude envelopes of the two networks are strongly coupled in terms of amplitude (**F**) and phase (**G**).

### Co-regulation of criticality and long-range communication through neuromodulation

Since neuromodulation can alter critical dynamics (Pfeffer et al., 2018), we tested whether it also affects the coupling and information transfer between networks. Neuromodulation was modelled as a multiplicative modulation in the synaptic strength of excitatory to excitatory connections (Fig. 5A) in the second network (N2), during the time intervals of 200 to 400 seconds, and 600 to 800 seconds (Fig. 5B,C). A modulation by a factor of 0.75 decreases overall activity (Fig. 5B) and brings networks towards a more subcritical regime for the duration of neuromodulation (Fig. 5D), while the opposite is true for a modulation by a factor of 1.25 (Fig. 5C,E). These changes in criticality are accompanied by alterations in amplitude, phase coupling and information transfer, for the duration of neuromodulation: both sub- and super-critical networks can transfer information once they are brought towards a critical regime through appropriate changes in synaptic strength (Fig. 5F-K). On the other hand, network coupling and information transfer revert back to baseline once neuromodulation is turned off. These results indicate that neuromodulation can regulate the ability of networks to establish long-range communication while maintaining a fixed structure of excitatory and inhibitory connections.

**Figure 5.**
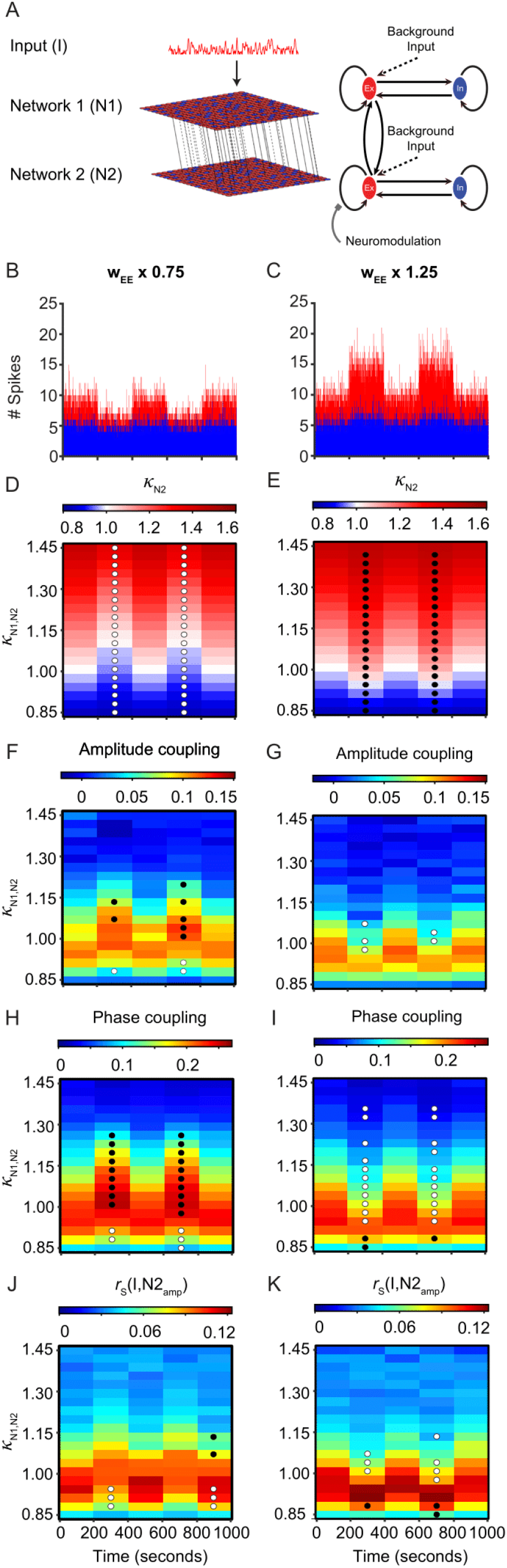
Neuromodulation of synaptic strength alters criticality with implications for network coupling and information processing. (**A**) Neuromodulation of activity in the second network is simulated by changing the synaptic strength of excitatory-to-excitatory connections. (**B**) Transient neuromodulation applied during time intervals of 200–400 seconds and 600–800 seconds leads to decreases in overall activity in case of multiplication by a sub-unitary factor (**B**, *w*_*EE*_ ∗ 0.75), and increases in activity for multiplication by a supra-unitary factor (**C**, *w*_*EE*_ ∗ 1.25). Consequently, the criticality of the networks also changes—making super-critical networks more critical for the sub-unitary multiplication factor (**D**), and sub-critical networks more critical for the supra-unitary multiplication factor (**E**). *Black/white* circles represent significant positive/negative change in statistics when compared to the baseline (significance threshold *p* ≤ 0.001). As neuromodulation changes the synaptic strength and with it the criticality of the networks, it also affects long-range communication. More specifically, when networks are brought away from criticality, amplitude coupling (**F,G**), phase coupling (**H,I**) and information transfer (**J,K)** are impaired. Shifting the networks closer to criticality by means of neuromodulation has the opposite effect, namely to facilitate long-range communication.

## Discussion

We have investigated how the local E/I ratio of neuronal networks shapes their spatio-temporal dynamics and capacity to communicate with each other through long-range connections. We observed that the long-range coupling of neuronal oscillations, as well as information transfer through oscillatory networks is greatly facilitated when networks operate close to the critical state, where excitation and inhibition are in balance. This relationship between criticality and network coupling is robust to changes in the number of long-range connections and is present for a wide range of conduction delays. In addition to the importance of the E/I ratio of synaptic connections, we show that neuromodulation of synaptic gain co-regulates critical dynamics and how networks couple to each other and transfer information. Thus, local criticality and E/I balance—as determined by both local connectivity and neuromodulatory gain—are generic mechanisms enabling long-range communication.

Our results confirmed our hypothesis that the critical state is conducive of better long-range coupling. The intuition behind this hypothesis lies in that the critical regime is characterized by transient synchrony, lying at the transition between a subcritical quiescent regime and a regime with strong, persistent oscillations (Dalla Porta and Copelli, 2019). In a subcritical regime, networks are not likely to exhibit long-range coupling: any input from an external network quickly dies out because of the strong inhibition. In the super-critical regime, on the contrary, networks are so active that any input coming from another network drowns in the relentless activity. It is only in the critical state, where localized activations have equal chance to spread or to die out, that the networks are able to phase and amplitude couple.

In our study, coupling is restricted to only two networks. However, in the brain, the constellation of areas that are functionally coupled is continuously changing (Deco et al., 2017; Tewarie et al., 2019). During rest, an intermediate inter-areal connection strength, which could correspond to excitation/inhibition balance with critical dynamics, enables the functional connectome to spontaneously reach the highest number of possible states (Deco et al., 2017). Our study suggests that, in addition to the strength of anatomical connections, local E/I balance and the associated critical dynamics are also important for flexible inter-areal coupling.

The high dimensionality of the state space in the critical regime is thought to allow functional networks to form and dissolve quickly in a task-dependent manner (Deco and Kringelbach, 2016). One possible mechanism for task dependent changes in coupling is demonstrated by Palmigiano et al. (2017), where a constant input attached to one of two networks, selectively increases phase coupling and biases information transfer. In our computational model, an external input altered the criticality of the receiving network making it more supercritical. Importantly, information transfer of the external input was maximal when the second, coupled network was critical. Such a pairing of a super-critical to a critical network leads to stronger phase coupling than if both networks are critical, possibly because the super-critical network exhibits a strong oscillatory regime with the power to entrain the oscillatory phase of the susceptible critical network.

The mechanisms by which the brain regulates the criticality of oscillations in the context of long-range coupling remain understudied. Neuromodulation, e.g., by locus coeruleus, could alter the coupling gain, with effects on criticality (Pfeffer et al., 2018), and as shown in our study, also on network coupling and information transfer. Another way in which the brain can influence long-range coupling is by shifting the inter-areal connectivity strengths, for example under the direction of the thalamus (Nakajima and Halassa, 2017). Three additional networks have been identified, with different modulatory effects on alpha oscillations (Sadaghiani and Kleinschmidt, 2016): the cingulo-opercular, fronto-parietal and dorsal attention networks. While cingulo-opercular and dorsal attention networks set the background level of alpha oscillations, it is the fronto-parietal network that establishes long-range phase coupling, and which is most involved in flexible cognitive control, e.g., in shaping the functional connectome during task switching (Gold et al., 2010; Marek and Dosenbach, 2018). However, coupling within the fronto-parietal network is altered in disorders like Alzheimer’s disease and schizophrenia (Babiloni et al., 2004; Sheffield et al., 2015). In the light of our findings, these alterations in long-range coupling could be explained by a departure from criticality, since evidence for locally more sub-critical alpha/beta oscillations have been reported for these disorders compared to healthy controls (Montez et al., 2009; Nikulin et al., 2012). Additionally, structural connectivity is damaged in these disorders (Bozzali et al., 2002; Davis et al., 2003), which could also impact network coupling. This is consistent with our model, where a reduction in the number of long-range connections leads to impaired amplitude- and phase-coupling, regardless of local criticality.

Phase and amplitude coupling represent distinct mechanisms (Siems and Siegel, 2020), which operate on different timescales (Daffertshofer et al., 2018): amplitude coupling is more persistent, whereas phase coupling is fast changing. However, there is also some degree of overlap between the underlying networks (Zhigalov et al., 2017; Siems and Siegel, 2020): in source-reconstructed MEG it was found that the amplitude, phase connectomes are co-localized with the criticality connectomes (Zhigalov et al., 2017). While this provides correlational evidence for a link between the two forms of long-range coupling and criticality, our study shows that local criticality is actually a pre-requisite for long-range coupling, as well as the ensuing information transfer.

Criticality in non-oscillatory systems has been associated with optimized information transmission in empirical studies, albeit only at small spatial scales in the millimeter range (multi-electrode arrays). For example, Shew et al. found that the critical state maximizes information transmission from an external stimulus to a local network (Shew et al., 2011). Our research complements this result, showing that criticality is also important for communication between distant networks exhibiting critical fluctuations in oscillatory activity.

When coupling two networks with the same level of local E/I ratio, we found that coupling super-critical to super-critical networks exhibit stronger phase coupling than critical-critical networks. Yang et al. reported similar results where local synchrony was found to be stronger when the local dynamics were in a super-critical regime (Yang et al., 2012). However, in the same study, the variability of synchronization was found to be reduced in the super-critical state, compared to criticality. Thus the strong, stable phase coupling seen in the super-critical state might prevent networks from flexibly establishing coupling when needed.

Sleep is considered important for homeostatic regulation of synaptic connections in the brain (Tononi and Cirelli, 2003). At a network level, sleep has also been related to the maintenance of E/I balance, critical dynamics, and long-range synchrony. Prolonged sleep deprivation leads to super-critical dynamics (Meisel et al., 2013), reduced synchronization of slow oscillations (Bu et al., 2017), alterations in network connectivity (Ben Simon et al., 2017), as well as changes in E/I balance (Chellappa et al., 2016; Bridi et al., 2020). The cognitive deficits (Alhola and Polo-Kantola, 2007) that follow sleep deprivation might be explained by altered functional integration with impaired long-range coupling. E/I balance, under the homeostatic influence of sleep, might serve as the mechanism that bridges between regulation at the synapse level and the emergent behavior.

Previous research has focused on the role of connectome topography and long-range coupling strength for brain-wide information integration. Here, we show that local excitation/inhibition balance, as reflected in criticality, is a pre-requisite for long-range coupling. Pharmaceutical interventions that target local E/I balance (Bruining et al., 2020), may thus help to also restore coupling and its functional consequences.

## Methods

### CRitical OScillations (CROS) model

We used an adapted version of the model described in (Poil et al., 2012), with synaptic weights optimized for power-law avalanches and long-range temporal correlation. The model was implemented using the Brian2 simulator for spiking neural networks (Stimberg et al., 2014; RRID:SCR_002998). Specifically, we modeled networks of 75% excitatory and 25% inhibitory integrate-and-fire neurons arranged on a 50×50 open grid. Placing neurons on the grid using uniform sampling may result, by chance, in clusters of excitatory or inhibitory neurons. To avoid this, we first positioned inhibitory neurons on the grid according to Mitchell’s best candidate algorithm (Mitchell, 1991), which results in a more even spatial distribution. The remaining spaces were filled with excitatory neurons. Networks differ in their two connectivity parameters, *C*_*E*_ and *C*_*I*_, which are the percentage of other neurons within a local range (a circle with radius = 4 neurons centered on the presynaptic neuron) that each excitatory and each inhibitory neuron connects to, respectively. Connectivity parameters were set between 5–100% at 5% intervals, and 10 different networks were created for each combination of *C*_*E*_ and *C*_*I*_. Border neurons had fewer connections because these neurons had a lower number of neurons in their local range. Within local range, connection probability decreases exponentially with distance. More specifically, the probability, *P*, of a connection at a distance *r* was given by:

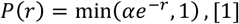

where *α* is optimized separately for excitatory and inhibitory neurons such that the overall connectivity probability within a neuron’s local range is equal to *C*_*E*_ or *C*_*I*_, depending on whether the neuron is excitatory or inhibitory. For example in the case of an excitatory neuron *i*, with connectivity probability *C*_*E*_, where the set of neighboring neurons within the local range is *J*, and |*J*| is the number of neighbors, *α* will have to satisfy the following equation:

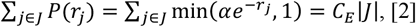

As such, we used the Nelder-Mead optimization algorithm to determine the value of *α* which minimizes the following function:

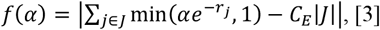

### Neuron model

Neurons were modeled using a synaptic model integrating received spikes, and a probabilistic spiking model. Each time step (*dt*) of 1 ms starts with each neuron, *i*, updating the input *I*_*i*_ with spikes received from the presynaptic neurons *J*_*i*_, together with an exponential synaptic decay:

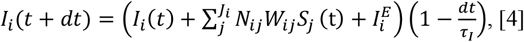

The weights *W*_*ij*_ are fixed, and depend on the type of the pre- and post-synaptic neuron. τ_*I*_ is the decay constant of inputs, and *S* is a is a binary vector, with *S*_*j*_=1 if the pre-synaptic neuron *j* fired in the previous time step, and *S*_*j*_=0 otherwise. 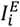 is the external input to the neurons which is 0 unless stated otherwise. *N*_*ij*_ is a multiplicative gain factor which is set to 1 unless stated otherwise

The activation of a neuron *A*_*i*_, is then updated with these excitatory and inhibitory inputs, together with an exponential decay, *τ_P_*, and a baseline activation level *A*_0_:

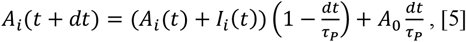

The spiking probability 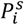 is calculated as a function of the neuron activation *A* at the current timestep, as follows:

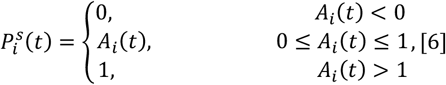

We determine whether the neuron spikes with the probability *P_S_*. If a neuron spikes, the neuron activation *A* is reset to the reset value, *A*_*r*_. At the next time step, all neurons that it connects to will have their input updated again according to Equation 4.

### Model parameters

All model parameters were the same as in the original paper (Poil et al., 2012) apart from the synaptic weights. Neuron model: (*τ*_*I*_ = 9 ms), Synaptic model: Excitatory neurons (*τ*_*P*_ = 6 ms, *A*_0_ = 0.000001, *A*_r_ = −2), Inhibitory neurons (*τ*_*P*_ = 12 ms, *A*_0_ =0, *A*_r_ = −20). To improve the range and stability of the long-range temporal correlations from the original model, an evolutionary algorithm (Smit and Eiben, 2011) was applied to the synaptic weights. The parameters that could vary were the 2 connectivity parameters (taking values between 0–100%), and the natural logarithm of the magnitude of the 4 synaptic weights, *W_EE_*, *W_IE_*, *W_EI_* and *W_II_* (taking values between −5 and 1). For each run, a fitness value was calculated based on the avalanches size and duration distributions (See Methods - Neuronal Avalanches), and the LRTCs (see Methods - Detrended fluctuation analysis of long-range temporal correlations).

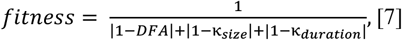

The optimum weights (*W_ij_*, connecting the presynaptic neuron *j* to the postsynaptic neuron *i*) found by the algorithm were (*W_EE_* = 0.0085,*W_IE_* = 0.0085,*W_EI_* = −0.569,*W_II_* = −2)

### Long-range connections

To assess how reciprocal long-range connections affect synchrony of oscillations and information transfer in our model, we generated two CROS model grids of 50×50 neurons. The placement of excitatory and inhibitory neurons on the two grids was exactly the same, to allow connection of neurons of the same type and at the same positions in the two networks. However, the local connectivity was allowed to vary between the two grids, according to the probabilistic connectivity rules described above. To create *N* long-range connections between the two networks (where *N* < number of excitatory neurons on the grid = 1875), we sample *N* excitatory positions. Half of these (*N*/2) are long-range connections from the first to the second network – projecting to excitatory neurons at the same positions on the grid. The other half connect from the second to the first network. This procedure ensures that there are no connections that go both ways.

### Conduction delay

0 ms for local connections within each grid. 25 ms on the long-range connections between the two grids, unless otherwise specified.

### Network input

A Gaussian white noise signal S was generated, with a length and sampling frequency equal to that of the network signal. The signal was then convoluted with an exponential: 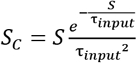 as in (Mainen and Sejnowski, 1995), where the parameter τ_*inputi*_ is the time constant of the signal, which allows to slow down the changes in the white noise signal. The absolute value was computed, |*S*_*C*_|, so that the input is only excitatory. This external input was connected to all the excitatory neurons of the first network. Thus, in equation [4], 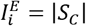.

### Network activity analysis

A network signal was created by summing the total number of neurons spiking at each time-step with a Gaussian white noise signal of the same length with mean = 0 and σ = 3. This level of white noise was set to allow all networks to achieve a time-varying phase, which is not the case without adding the noise, when there are silent periods in the network. Amplitude envelopes were extracted from this pre-processed signal, with Gaussian noise added. The analysis of neuronal avalanches was performed on the raw signal, consisting of the total number of neurons spiking at each time step.

### Oscillation power

The power spectrum was computed using the Welch method with a Hamming window of size with 2^11^ FFT points.

### Neuronal avalanches

A neuronal avalanche is defined as a period where neurons are spiking above a certain threshold—in our case set to half median of activity. The size of the avalanche is the number of spikes during this period. We then computed the κ index (Shew et al., 2009; Poil et al., 2012), which calculates the difference between the distribution of our data and a power-law, by calculating the average difference of the cumulative distribution of a power-law function, *P*, (with exponent −1.5 for size and −2.0 for duration) and that of our experimental data, *A*, at 10 equally spaced points on a logarithmic axis (*β*) and adding 1.

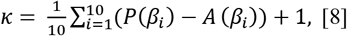

A subcritical distribution is characterized by *κ* < 1, and a super-critical distribution by *κ* > 1, whereas *κ* = 1 indicates a critical network.

### Neuronal oscillations

We focused our analysis of oscillations in the alpha band (8—16 Hz range), which was the most prominent oscillatory regime exhibited in our model (Poil et al., 2012).

### Detrended fluctuation analysis of long-range temporal correlations

Detrended fluctuation analysis (DFA) was used to analyze the scale-free decay of temporal (auto)correlations, also known as long-range temporal correlations (LRTC). The DFA was introduced as a method to quantify correlations in complex data with less strict assumptions about the stationarity of the signal than the classical autocorrelation function or power spectral density (Linkenkaer-Hansen et al., 2001; Hardstone et al., 2012). In addition, DFA facilitates a reliable analysis of LRTC up to time scales of at least 10% of the duration of the signal (Chen et al., 2002; Gao et al., 2006). DFA exponents in the interval of 0.5 to 1.0 indicate scale-free temporal correlations (autocorrelations), whereas an exponent of 0.5 characterizes an uncorrelated signal. The main steps from the broadband signal to the quantification of LRTC using DFA have been explained in detail previously (Linkenkaer-Hansen et al., 2001; Hardstone et al., 2012). In brief, the DFA measures the power-law scaling of the root-mean-square fluctuation of the integrated and linearly detrended signals, *F*(*t*), as a function of time-window size, *t* (with an overlap of 50% between windows). The DFA exponent is the slope of the fluctuation function *F*(*t*), and can be interpreted as the strength of the autocorrelations in signals. For our analyses we computed DFA on the amplitude envelope of the signal filtered in the 8–16 Hz range, and the exponent was fit between 5 and 30 seconds.

### Simulations

To determine the relation between connectivity and criticality, as well as phase- and amplitude-coupling, we set excitatory/inhibitory connectivity between 5—100% at 5% intervals, and 10 different network ensembles were created for each combination of *C*_*E*_ and *C*_*I*_. Both networks in each ensemble had the same local excitatory/inhibitory connectivity. We compared the results between *N* = 0 long-range connections (unconnected), and *N* = 400 long-range connections, with a conductance delay of 25 ms over the long-range connections. All simulations were run for 1000 seconds.

We then picked a diagonal across the critical line, where we varied excitatory connectivity between 5% and 100%, in steps of 2%. Inhibitory connectivity was computed as 105% minus excitatory connectivity. Additionally we varied the number of long-range connections (between 100 and 1800 in steps of 100) for all networks across the diagonal, 5 runs per parameter combination. Separately, we also varied conductance delays (between 0 and 100 ms, in steps of 5 ms)) for all networks across the diagonal, 5 runs per parameter combination. To analyze the effects of varying E/I balance between the two networks, we ran all combinations in one diagonal (first network) against all combinations in one diagonal (second network), 5 runs for each parameter combination. The same approach was used with the external input, where, in addition to conduction delays and the number of long-range connections, we also varied the time constant of the input signal, τ_*input*_, where we sampled 15 values on a log scale between 3 and 800 ms. In the case of neuromodulation, we started from the same diagonal across the critical line, and ran simulations for two levels of synaptic gain (0.75 and 1.25). Throughout the 1000 seconds of each simulation, the synaptic gain *N*_*ij*_ of excitatory to excitatory connections (equation [4]) alternated between the baseline level (*N*_*ij*_ = 1) and another value (0.75 or 1.25), every 200 seconds. For each 200 seconds interval where the synaptic gain was is maintained constant, we computed criticality (DFA and κ), as well as network coupling and information transfer. Since the reliability of estimates is lower for these shorter (200 seconds) intervals compared to the same estimates computed for the entire 1000 seconds of the simulations, we ran 20 networks for each parameter combination.

### Amplitude coupling

Amplitude coupling between the two networks was measured as the Spearman correlation between the amplitude envelopes of the signals filtered in the 8—16 Hz range. This value was compared with a surrogate where the causal relationship between the two signals was destroyed by swapping the two halves of one of the signals.

### Phase coupling

Phase coupling was calculated using inter-site phase clustering (Lachaux et al., 1999; Cohen, 2014). The advantage of this method, compared to magnitude-squared coherence, is that it does not include information about oscillation amplitude, and thereby allows us to isolate amplitude from phase coupling. First we computed the amplitude envelope of the signals *x* and *y*, filtered in the 8–16 Hz range. Then we extracted the phase by taking the angle of the Hilbert transform *ϕ*_*x*_, *ϕ*_*y*_. Last, we computed the uniformity of phases differences across all time points. For this calculation, we used the circ_r function from the circ_stats package (Berens, 2009):

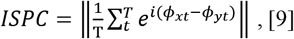

Phase coupling was then compared with a surrogate where the causal relationship between the two signals was destroyed by swapping the two halves of one of the signals.

### Information transfer

We correlated the input signal with the amplitude envelope of alpha oscillations (8–16 Hz) of the first network *r*_*S*_(*I*, *N*_1_) and the second network *r*_*S*_(*I*, *N*_2_). Since the input has to cross network 1 to get to network 2, we took *r*_*S*_(*I*, *N*_2_) as a measure of information transfer. Specifically, we computed *r*_*S*_(*I*, *N*_2_) at different time lags (between 0 and 500 ms), and took the maximal value to be the information transfer. Information transfer was then compared with a surrogate where the causal relationship between the input and the two signals was destroyed by swapping the two halves of one of the signals. To calculate the lag at which the input arrives in the second network, we determined the time point where *r*_*S*_(*I*, *N*_2_) is maximal (between 0 and 500 ms). Since *r*_*S*_(*I*, *N*_2_) can show random fluctuations around zero even when no information transfer occurs (e.g. in unconnected networks), we considered time lags only for coupled networks where *r*_*S*_(*I*, *N*_2_) was greater than a fixed threshold = 0.04.

## ACKNOWLEDGMENTS

This work was funded by Netherlands Organization for Scientific Research (NWO) Social Sciences grant 406.15.256 (to K.L.-H. and A.-E.A.). Also supported by the program committee ‘Systems and Network neuroscience’ of Amsterdam Neuroscience (to K.L.-H.).

## Supplementary information

**Figure S1.**
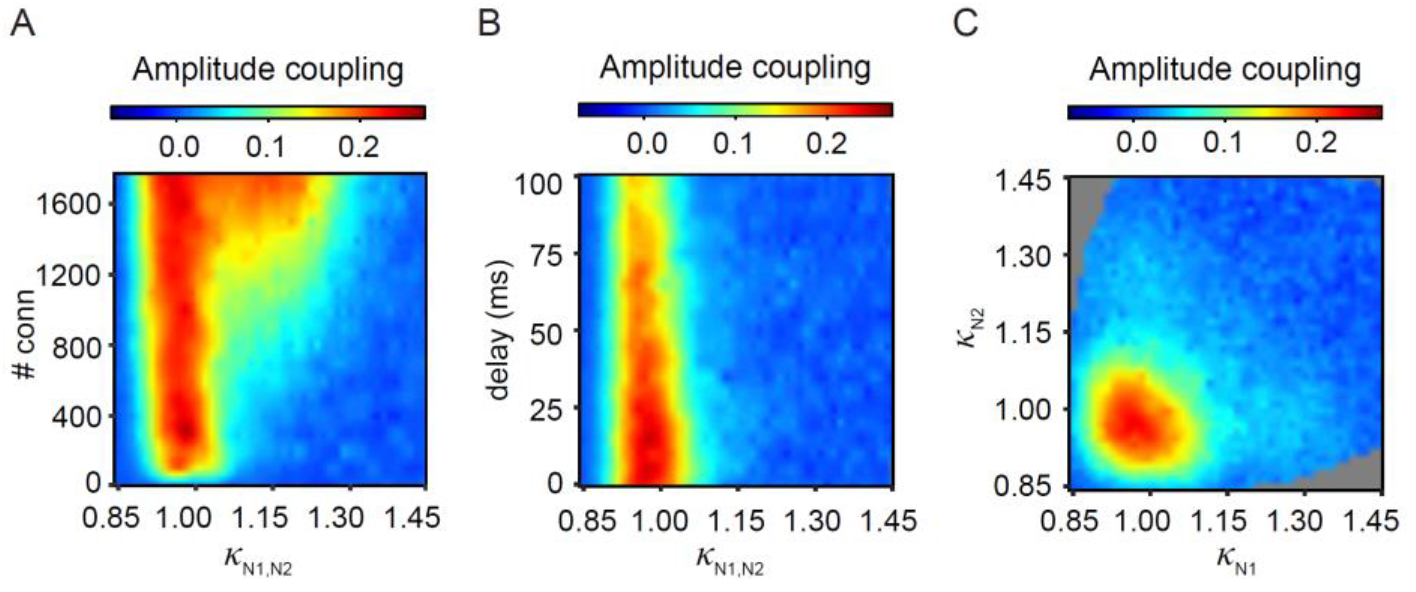
Maximal amplitude correlation at criticality regardless of parameter changes. **(A)** The correlation of amplitude envelopes close to criticality is robust to increases in the number of long-range connections, as to increases in conduction delay of long-range connections (**B**). If the number of long-range connections is very large (>1200), super-critical networks can also show coupling of amplitude envelopes of alpha oscillations (**A**). (**C**) We varied the level of criticality between the two networks, e.g., allowing for coupling of super-critical-subcritical networks. The number of connections was kept at 400, and the network combinations were chosen from the diagonal in (**Fig. 2B**). Amplitude envelopes are coupled only when both networks are close to criticality.

**Figure S2.**
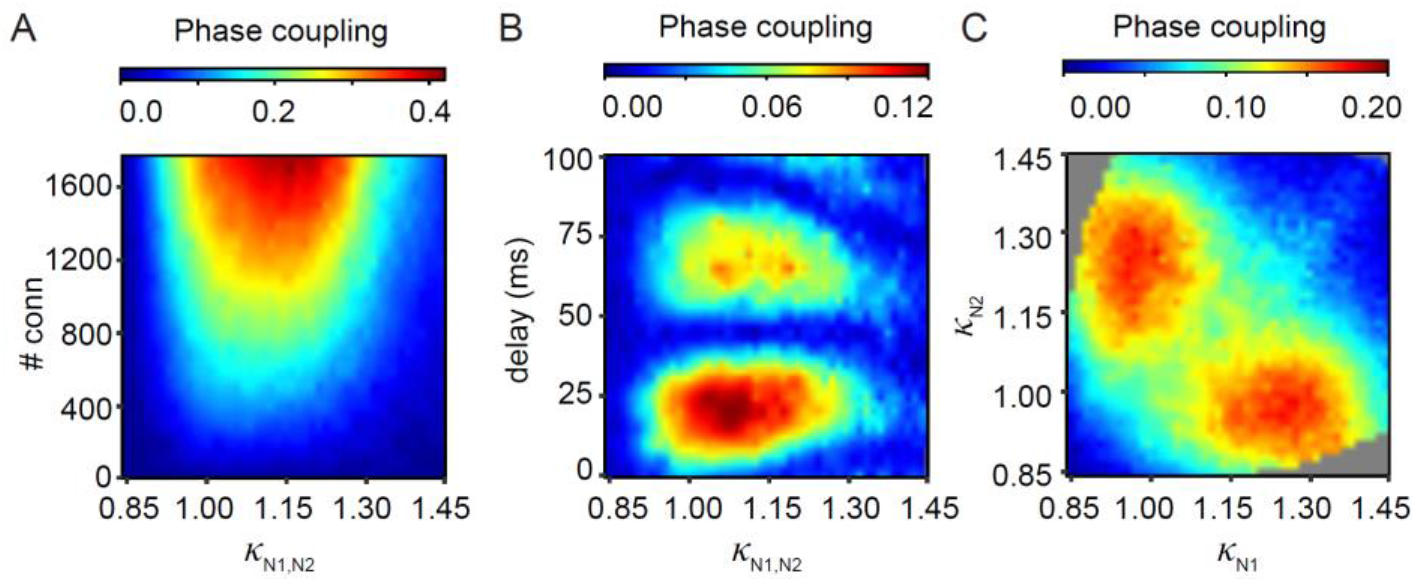
Influence of model parameters on phase coupling. **(A)** Phase coupling increases with the number of long-range connections between the two networks. **(B)** Phase coupling emerges at specific conduction delays (*N* = 400 long-range connections). **(C)** Varying E/I balance of the two networks independently of each other reveals that maximal phase coupling occurs when one network is close to criticality (*κ* ~ 1), and the other is super-critical (*κ* ~ 1.2).

**Figure S3.**
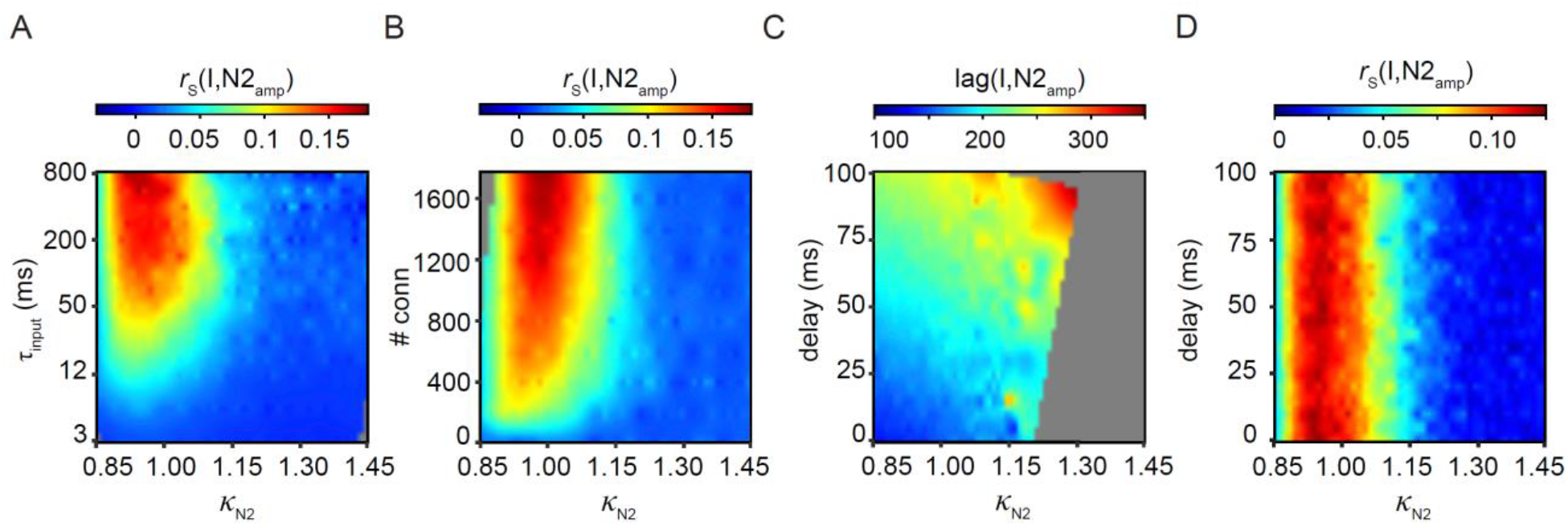
Influence of model parameters on information transfer. Stimulus coupling is maximal when the second network is critical 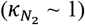 also when changing the timescale of the input signal (**A**) or the number of connections (**B**). These parameters ultimately affect stimulus coupling, which increases for signals with longer timescales (**A**) and networks coupled with more long-range connections (**B**). Last, we manipulated the conductance delay between the two networks. At higher conductance delays, it took longer for the input to reach N2, which was reflected in the time lag at which maximum correlation with the input was achieved (**C**). However, at its peak, the correlation with the input was unaffected by changes in conductance delays (**D**).

